# Age-Associated Behavioral Alterations in Laboratory-Housed Octodon degus

**DOI:** 10.64898/2026.07.07.737045

**Authors:** Huiru Bai, Yiran Liu, Andrei Seluanov, Vera Gorbunova

## Abstract

Octodon degus are long-lived, diurnal, and highly social rodents increasingly used in studies of aging and neurodegeneration. However, behavioral profiles of aged laboratory-housed degus remain incompletely characterized, limiting the interpretation of aging-associated functional and molecular phenotypes in this species. Here, we evaluated age-associated changes in locomotor activity, open-field exploration, social novelty behavior, and manually scored ethological responses in young and old degus maintained under long-term laboratory housing conditions. Automated behavioral tracking was performed during open-field testing and three-chamber social behavior testing. During open-field testing, old degus showed increased locomotor activity compared with young animals, including greater total distance moved, higher mean velocity, increased moving frequency, and longer cumulative movement duration. Old degus also showed increased center-zone duration and reduced thigmotaxis score. Manual ethological scoring revealed increased rearing and fecal boli in old animals during open-field exposure. In the three-chamber social behavior assay, young degus showed higher investigation frequency toward the novel intruder than toward the familiar cagemate, whereas old degus showed a lower social novelty discrimination index compared with young animals. Sex-stratified analyses did not identify significant male–female differences within young or old groups for the major open-field or social novelty metrics examined. Together, these findings indicate that aging in laboratory-housed degus is associated with a mixed behavioral profile involving increased stress-related ethological responses and reduced social novelty preference reminiscent of dementia-like behavioral changes observed in Alzheimer’s disease. This behavioral framework provides a practical reference for future studies examining behavioral heterogeneity and molecular correlates of brain aging in degus.

**Lay Summary:** Octodon degus are long-lived, highly social rodents that are increasingly used to study aging and age-related neurodegenerative disorders. However, interpreting behavioral changes in aged degus requires a clear understanding of how aging affects activity, exploration, social behavior, and stress-related responses under laboratory housing conditions. In this study, we compared young and old degus using open-field testing, three-chamber social behavior testing, automated video tracking, and manual scoring of selected behaviors. Aged degus did not show a simple reduction in behavioral activity. Instead, they showed increased movement during behavioral testing, increased rearing behavior, greater exploration of the center of the open-field arena, and increased fecal output during open-field exposure. These findings suggest that aged degus show increased exploratory activity together with altered stress-related responses in a novel environment. Aged degus also showed reduced preference for investigating a novel social partner, consistent with a dementia-like cognitive impairment. Together, these results define a behavioral profile of aged laboratory-housed degus and provide a practical reference for future studies using this species to investigate aging, social behavior, and neurodegeneration-related phenotypes.

## 1. Introduction

Octodon degus are long-lived, diurnal, and highly social hystricomorph rodents that have become increasingly important in studies of aging and age-related neurodegenerative disease. Degus are of particular interest because aged animals can develop behavioral impairment and Alzheimer’s disease (AD)-like neuropathological features without genetic engineering, including amyloid-β accumulation, tau-related pathology, synaptic dysfunction, neuroinflammation, and cognitive decline in subsets of aged animals^1–8^. Unlike transgenic mouse models that reproduce selected familial AD-related molecular features, degus provide a natural aging context in which behavioral and neuropathological phenotypes emerge heterogeneously across animals and colonies^4,9–14^. This heterogeneity resembles an important feature of natural aging, but it also creates a need for careful behavioral characterization under defined laboratory conditions.

Behavioral phenotyping is essential for interpreting brain aging and neurodegenerative vulnerability. In humans and animal models, neurodegenerative disease is not limited to memory impairment; it can also involve changes in locomotor activity, exploratory drive, anxiety- or stress-related behavior, arousal regulation, social interaction, and novelty recognition^15–18^. These domains are especially relevant for degus, a highly social species that is responsive to environmental novelty and has been used to model age-associated cognitive and AD-like phenotypes^1,2^. Therefore, behavioral measures such as open-field exploration, thigmotaxis, social novelty preference, and ethological responses may provide useful functional readouts of neural aging in this species.

Previous degu studies have used several behavioral paradigms to evaluate age-associated cognitive and functional decline, including burrowing behavior, social interaction, novel object and novel location recognition, and Barnes maze learning^4,19,20^. Burrowing has been particularly influential because it represents an ethologically relevant, activity-of-daily-living-like behavior; impaired burrowing performance has been reported in aged degus and has been associated with cognitive impairment and AD-like neuropathological features^21^. Other studies have identified age-associated deficits in recognition memory, spatial learning, and social behavior, further supporting the value of degus for studying behavioral aspects of neurodegenerative aging^4,19,20^.

Despite these advances, several behaviorally important domains remain incompletely integrated in aged laboratory-housed degus. In standard rodent studies, open-field testing is widely used to assess locomotor activity, exploratory behavior, and anxiety-related center–border behavior, including thigmotaxis^22^. Social interaction and three-chamber assays are commonly used to assess sociability, social investigation, and social novelty preference^23–25^. Manual ethological scoring can further capture behaviors such as rearing, grooming, jumping, and fecal output, which provide additional information about exploration, arousal, and stress-related responses^22,26–29^. However, age-associated changes in locomotor activation, thigmotaxis, social novelty preference, and open-field ethological responses have not been systematically examined together in the same laboratory-housed degu aging cohort. This gap limits the interpretation of behavioral phenotypes in aged degus because changes in activity, exploration, stress responsiveness, and social novelty can each influence the interpretation of cognitive and neurodegeneration-associated outcomes.

Here, we characterized age-associated changes in locomotor activity, open-field exploration, social novelty behavior, and manually scored ethological responses in young and old laboratory-housed Octodon degus maintained under long-term colony conditions. By integrating automated behavioral tracking with manual ethological scoring, we tested whether aging in degus is associated with generalized behavioral suppression or with domain-specific changes across exploratory, stress-related, and social behavioral measures. This study identifies age-associated behavioral alterations relevant to neurodegenerative aging and provides a functional framework for future studies linking degu behavior to molecular and neuropathological features of brain aging.

## 2. Materials and Methods

### 2.1. Animals

Octodon degus were maintained in the laboratory colony at the University of Rochester. Animals were grouped by age into young and old cohorts. Young animals were defined as <2 years of age, and old animals were defined as >4 years of age. Both male and female animals were included in the study. Assay-specific sample sizes are provided in Figure 1B. Briefly, open-field locomotor tracking included 19 young degus and 31 old degus, chamber-based locomotor tracking included 15 young degus and 26 old degus, and manual ethological scoring was performed using open-field recordings from 19 young degus and 31 old degus.

All animal procedures were performed in accordance with institutional guidelines and were approved by the University Committee on Animal Resources at the University of Rochester under protocol UCAR-2009-054R.

### 2.2. Housing and Husbandry

Degus were maintained under long-term laboratory housing conditions with group housing, standardized diet, bedding, and environmental enrichment^30,31^. Two primary cage systems were used for adult colony housing. The original cage system consisted of Tecniplast Static Cage 2000P units with 320 in² of floor space, including a cage base, wire top, and food hopper (Tecniplast, Tecniplast Static Cage 320 in²-2000P, 2000P001). These cages housed up to three adult degus and could accommodate an exercise wheel. A larger cage system consisted of Tecniplast GP-Suite guinea pig cages, each providing 618 in² of floor space and measuring approximately 31.875 × 28.000 × 9.875 inches (Tecniplast, GP-SUITE - Rack for Guinea Pigs). These cages housed up to five adult degus and allowed placement of both an exercise wheel and a shelter.

Biofresh Comfort Bedding (ScottPharma Solutions, L0110 Biofresh Comfort Bedding Natural 60 liter) was used as bedding, and each cage was provided with an Enviropak (ScottPharma Solutions, EnviroPak). Environmental enrichment included Exotic Nutrition exercise wheels (Exotic Nutrition Pet Supply, Treadmill Wheel), wooden huts or wooden cubes for chewing, stainless steel toys, and weekly dust baths. Dust baths were provided using Blue Beauty Dust placed in a stainless-steel bowl, typically the day before cage changes.

Degus were fed LabDiet 5025 Guinea Pig Diet (LabDiet, 5025 - Guinea Pig Diet) daily using cage food hoppers. Plain tap water was provided ad libitum in water bottles. Animals also had continuous access to Timothy hay cubes and received small amounts of whole oats sprinkled into the bedding several times per week, particularly during cage changes, to encourage foraging behavior. Animals were monitored as part of routine colony husbandry and veterinary care procedures.

### 2.3. Open-Field Behavioral Testing

Open-field behavior was assessed in a square arena measuring 100 cm × 100 cm with 40 cm-high grey walls and a white floor. This color configuration provided visual contrast between the animal and the arena background for automated video tracking. Each animal was placed individually into the arena and allowed to explore freely for 5 min. Behavioral videos were recorded from above the arena and analyzed with EthoVision XT 16 software (Noldus Information Technology, Wageningen, The Netherlands). The arena was cleaned between animals to reduce odor cues.

For center–border analysis, the arena was divided into a central square zone measuring 50 cm × 50 cm and a surrounding border zone. Because the full arena measured 100 cm × 100 cm, the center zone represented 25% of the total arena area and the border zone represented 75% of the total arena area.

### 2.4. Three-Chamber Social Behavior Testing

Social behavior was assessed using a three-chamber apparatus measuring 120 cm in total length and consisting of three 40 cm chambers connected by openings between adjacent chambers. The assay consisted of three sequential 10-min phases: habituation, sociability, and social novelty. During habituation, the test animal was allowed to freely explore the empty apparatus. During the sociability phase, a familiar cagemate was placed inside a wire enclosure in one side chamber, while the opposite side chamber contained an empty wire enclosure. During the social novelty phase, a novel intruder animal was placed inside the previously empty wire enclosure, while the familiar cagemate remained in the opposite side chamber. The side containing the cagemate or intruder was counterbalanced when possible. Behavioral videos were recorded from above and analyzed using EthoVision XT 16.

### 2.5. Manual Ethological Scoring

To complement automated locomotor tracking, selected ethological behaviors were manually scored from open-field recordings over the same 5-min observation period. The scored behaviors included fecal boli, rearing events, grooming events, and jumping events. Fecal boli were quantified as the total number of fecal pellets produced during the scoring window. Rearing was scored when the animal raised its forelimbs from the floor in a vertical exploratory posture. Grooming was scored as visible self-grooming behavior. Jumping was scored when the animal lifted all four paws from the floor in a vertical jumping movement. Manual scoring was performed from recorded videos using the same predefined behavioral criteria for all animals, and scored values were analyzed as event counts over the 5-min open-field exposure.

### 2.6. Behavioral Tracking and Data Processing

Behavioral videos were analyzed using EthoVision XT 16. For automated tracking, the center-point position of each animal was tracked throughout each recording. Tracking quality was visually inspected before downstream analysis, and recordings were excluded from a given analysis if the animal did not complete the assay, if the video quality was insufficient for reliable tracking or manual scoring, or if the animal could not be consistently detected by the software. Because assay completion and recording quality differed across behavioral tests, sample sizes were reported separately for each assay and analysis.

For open-field analysis, automated tracking metrics included total distance moved, mean velocity, moving frequency, cumulative movement duration, center-zone cumulative duration, and area-corrected thigmotaxis score. Moving frequency was defined as the number of movement bouts detected during the recording window, and cumulative movement duration was defined as the total time classified as moving by the tracking software. Center-zone cumulative duration was quantified from center-point tracking data. The area-corrected thigmotaxis score was calculated as the percentage of time spent in the border zone minus the expected border-zone area proportion of 75%. Thus, positive values indicate greater-than-expected border-zone occupancy, whereas values closer to zero indicate border-zone occupancy closer to the area-based expectation.

For chamber-based locomotor analysis, total distance moved and mean velocity were calculated over the 10-min tracking window. These locomotor metrics were analyzed within each assay. Raw locomotor values were not directly compared across assays with different recording durations or apparatus configurations.

For three-chamber social behavior analysis, sniffing zones were defined as regions extending 5 cm around each wire enclosure. Sniffing-zone cumulative duration and sniffing-zone investigation frequency were quantified for the relevant zones in each phase. During the habituation phase, sniffing-zone preference was calculated using the two empty enclosure-associated sniffing zones to assess baseline zone bias before the introduction of social stimuli. During the sociability phase, the sociability discrimination index was calculated as (cagemate − empty enclosure) / (cagemate + empty enclosure) using sniffing-zone cumulative duration. During the social novelty phase, the social novelty discrimination index was calculated as (novel intruder − familiar cagemate) / (novel intruder + familiar cagemate) using sniffing-zone cumulative duration. Animals with zero cumulative duration in both zones used for a given discrimination index had an undefined index and were excluded from that index-based analysis.

### 2.7. Statistical Analysis

Statistical analyses were performed using GraphPad Prism 10. Data are presented as mean ± standard deviation (SD) with individual animals overlaid. Comparisons between young and old animals were performed using two-tailed Mann–Whitney tests. For comparisons between cagemate and intruder sniffing-zone investigation frequency within the same animals, paired two-tailed t tests were used. For sex-stratified analyses, males and females were compared within each age group using Šídák’s multiple comparisons test. Animals with zero cumulative duration in both the cagemate and intruder sniffing zones had an undefined social novelty discrimination index and were excluded from discrimination index analysis. A p value < 0.05 was considered statistically significant. Exact p values are shown in the corresponding figures.

## 3. Results

### 3.1. Study design, cohort structure, and behavioral workflow

To establish a baseline behavioral framework for aged laboratory-housed Octodon degus, we analyzed locomotor activity, open-field spatial use, social behavior, and manually scored ethological behaviors in young and old animals. Young animals were defined as <2 years of age, and old animals were defined as >4 years of age (Figure 1B). Because assay completion and recording quality varied across behavioral tests, the number of animals included in each analysis was reported separately. Open-field testing included 19 young degus and 31 old degus, three-chamber social novelty testing included 15 young degus and 26 old degus, and manual ethological scoring was performed using open-field recordings from 19 young degus and 31 old degus, with male and female sample sizes indicated in Figure 1B.

Animals were maintained under long-term laboratory housing conditions with group housing, standardized diet, and environmental enrichment. The experimental workflow consisted of open-field testing, three-chamber social novelty testing, automated behavioral tracking, and manual ethological scoring. Open-field tracking was used to quantify locomotor activity and spatial-use metrics, including total distance moved, mean velocity, moving frequency, cumulative movement duration, center-zone duration, and thigmotaxis score. Social behavior analyses included sociability, social novelty, sniffing-zone investigation frequency, and social novelty discrimination index. Manual ethological scoring was used to assess fecal boli, rearing, grooming, and jumping during open-field exposure (Figure 1A). This design allowed us to assess age-associated changes across locomotor, spatial-use, social, and manually scored ethological behavioral domains.

**Figure 1.**
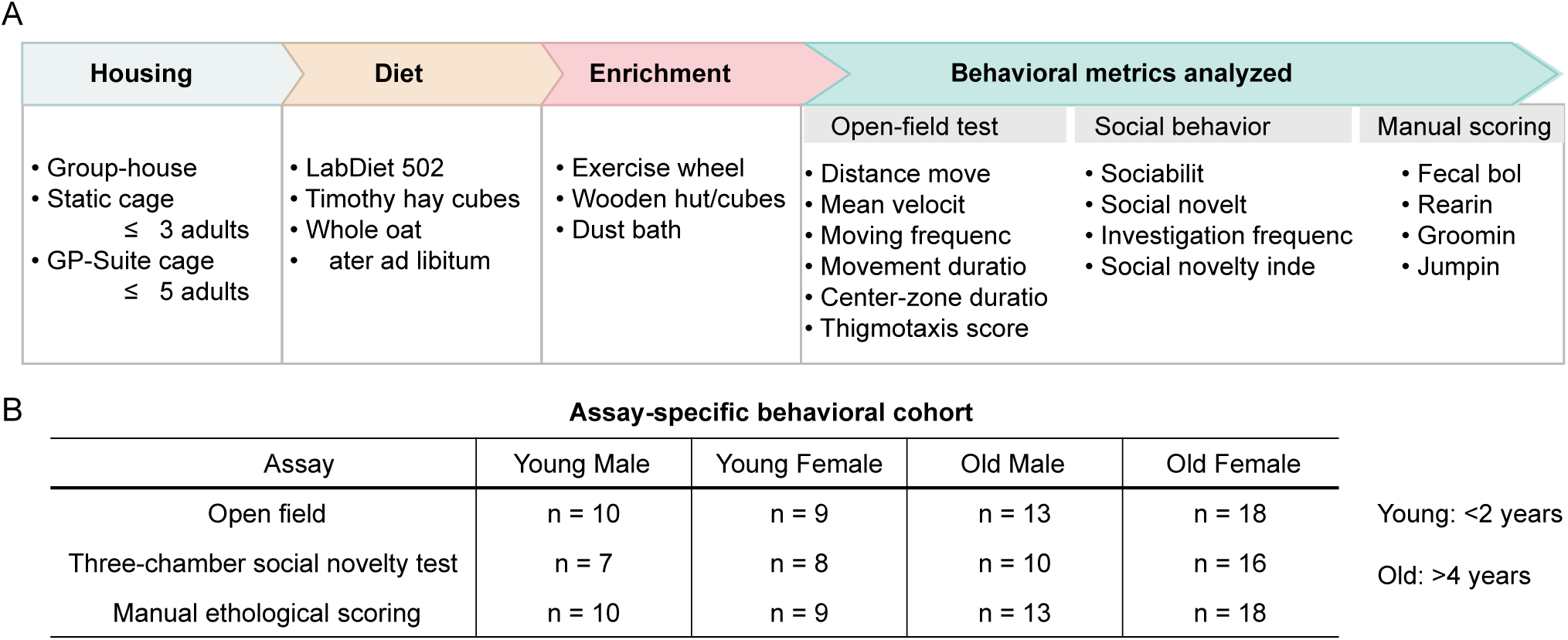
Study design and behavioral analysis overview. (A) Schematic overview of laboratory housing and husbandry conditions, behavioral workflow, and analyzed behavioral metrics. Open-field analyses included locomotor tracking and spatial-use measures during a 5-min recording window. Three-chamber social behavior testing included three sequential 10-min phases: habituation, sociability, and social novelty. Sniffing-zone investigation frequency and social novelty discrimination index were quantified during the social novelty phase. Manual ethological scoring was performed over a 5-min open-field exposure and included fecal boli, rearing, grooming, and jumping. (B) Assay-specific cohort structure for open-field testing, three-chamber social novelty testing, and manual ethological scoring. Young animals were defined as <2 years of age, and old animals were defined as >4 years of age. Values indicate the number of animals included in each analysis.

### 3.2. Aged degus show increased locomotor activity and reduced thigmotaxis in the open-field arena

We first assessed age-associated locomotor activity and center–border exploration during a 5-min open-field tracking window. Representative locomotor trajectories and spatial activity heatmaps showed broader arena exploration and greater activity density in old degus compared with young animals in both males and females (Figure 2A – E). Quantitative analysis confirmed that old degus moved a significantly greater total distance than young degus (p = 0.0004; Figure 2F). Old degus also exhibited higher mean velocity (p = 0.0009; Figure 2G), increased moving frequency (p = 0.0002; Figure 2H), and longer cumulative movement duration (p = 0.0003; Figure 2I).

We next quantified center-zone duration and area-corrected thigmotaxis score to assess center–border exploration. Old degus showed increased center-zone duration (p = 0.0003; Figure 2J) and a reduced area-corrected thigmotaxis score (p = 0.0004; Figure 2K). Together, these findings indicate that aged degus displayed increased locomotor activity together with reduced thigmotaxis and increased center-zone exploration during open-field exposure.

**Figure 2.**
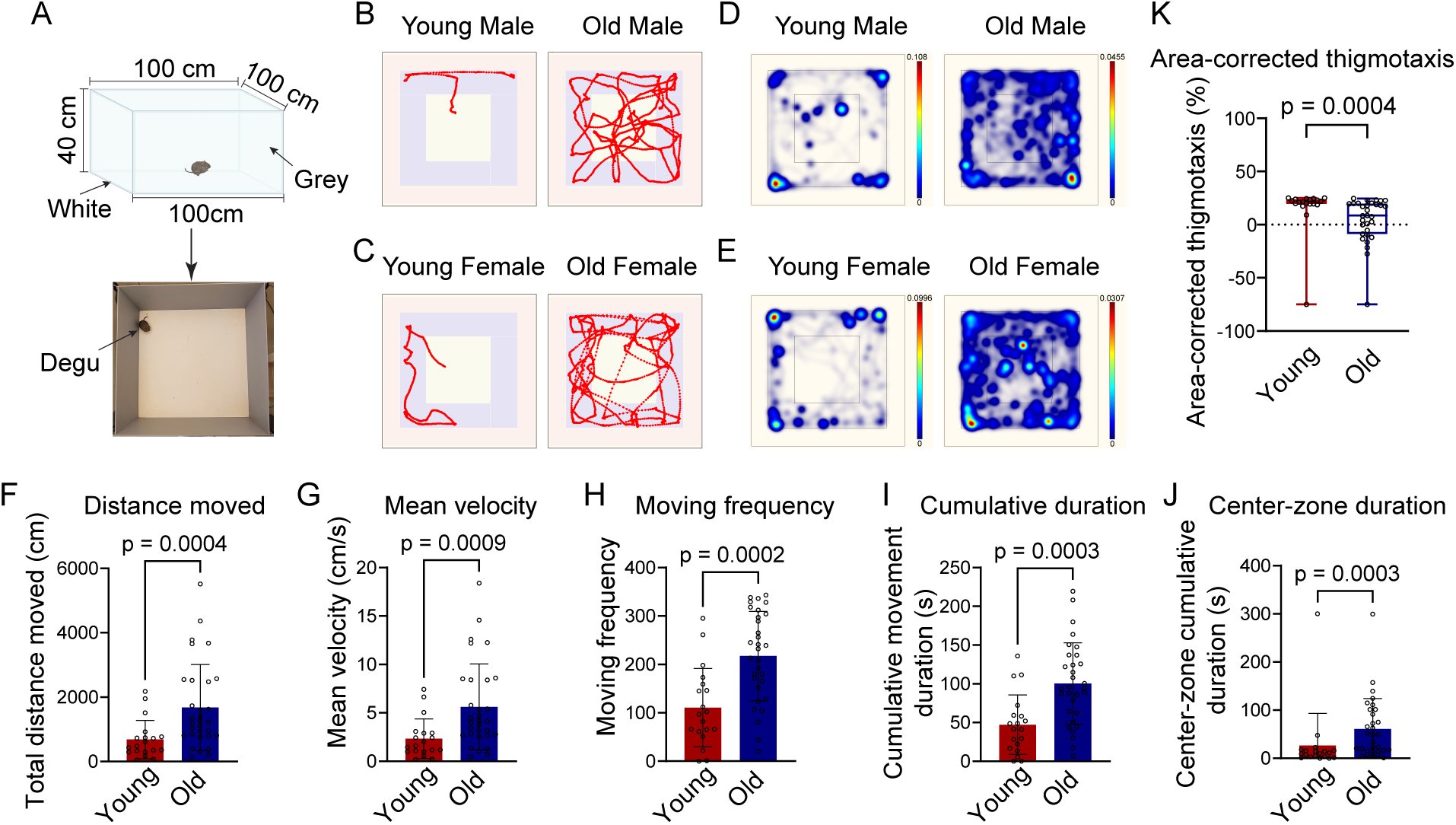
Open-field locomotor activity and spatial-use patterns in young and old degus. (A) Schematic and representative image of the open-field arena used for behavioral testing. (B, C) Representative locomotor trajectories from young and old male (B) and female (C) degus during the 5-min open-field tracking window. (D, E) Representative spatial activity heatmaps from young and old male (D) and female (E) degus. (F – I) Quantification of automated open-field locomotor metrics, including total distance moved (F), mean velocity (G), moving frequency (H), and cumulative movement duration (I). (J, K) Quantification of center-zone cumulative duration (J) and area-corrected thigmotaxis score (K) as exploratory spatial-use measures. The area-corrected thigmotaxis score was calculated as the percentage of time spent in the border zone minus the expected border-zone area proportion of 75%. Data are presented as mean ± SD with individual animals overlaid; n = 19 young degus (10 males and 9 females) and 31 old degus (13 males and 18 females). Statistical significance was determined using a two-tailed Mann–Whitney test.

### 3.3. Aged degus show increased locomotor activity in a chamber-based behavioral apparatus

We next asked whether the age-associated increase in locomotor activity observed in the open-field arena was also present in a structured chamber-based behavioral environment. Movement was analyzed during a 10-min chamber-based tracking window (Figure 3A). Representative locomotor trajectories and spatial activity heatmaps showed greater movement throughout the apparatus in old degus compared with young animals in both males and females (Figure 3B – E). Quantitative analysis confirmed that old degus moved a significantly greater total distance than young degus (p = 0.0046; Figure 3F) and showed higher mean velocity during chamber-based testing (p = 0.0004; Figure 3G). Together, these findings indicate that increased locomotor activity in aged degus was not limited to the open-field arena but was also detected in a structured chamber-based behavioral environment.

**Figure 3.**
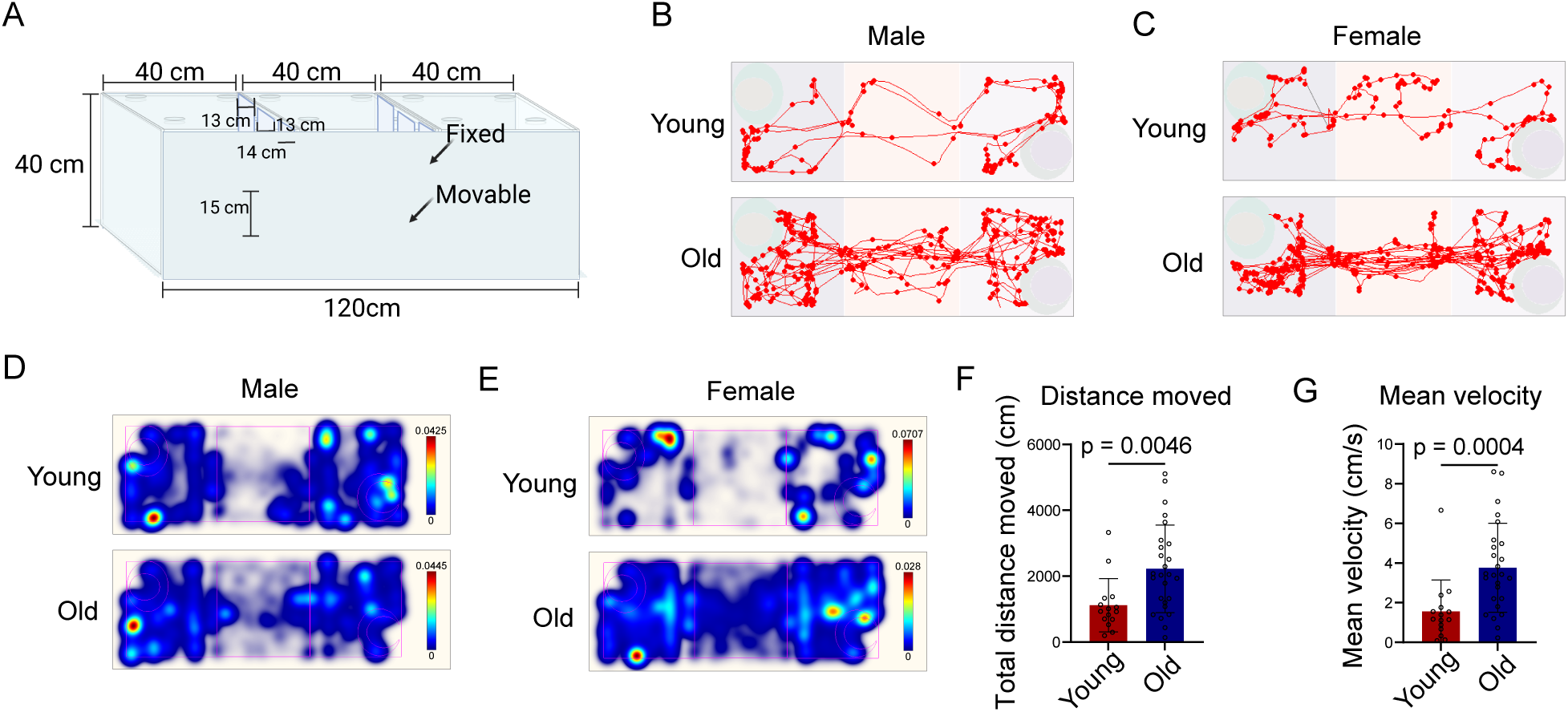
Chamber-based locomotor activity is increased in aged degus. (A) Schematic of the chamber-based behavioral apparatus. (B, C) Representative locomotor trajectories from young and old male (B) and female (C) degus during the 10-min chamber-based tracking window. (D, E) Representative spatial activity heatmaps from young and old male (D) and female (E) degus. (F, G) Quantification of total distance moved (F) and mean velocity (G) during chamber-based behavioral testing. Old degus showed significantly higher total distance moved and mean velocity than young degus. Data are presented as mean ± SD with individual animals overlaid; young, n = 15 (7 males and 8 females); old, n = 26 (10 males and 16 females). Statistical significance was determined using a two-tailed Mann–Whitney test.

### 3.4. Aged degus show reduced social novelty preference

We next assessed social behavior using a three-chamber social behavior assay consisting of habituation, sociability, and social novelty phases. Representative locomotor trajectories were visualized across all three phases in young and old degus (Figure 4A). To evaluate potential baseline preference for the sniffing zones before introduction of social stimuli, we first quantified sniffing-zone preference during the habituation phase. Young and old degus did not differ significantly in baseline sniffing-zone preference index (p = 0.3688; Figure 4B), suggesting that age-associated differences during the social novelty phase were unlikely to be explained by pre-existing sniffing-zone bias.

We next assessed sociability during the sociability phase. The sociability discrimination index did not differ significantly between young and old degus (p = 0.0862; Figure 4C), indicating that aged animals did not show a significant loss of general social approach toward a familiar cagemate under these testing conditions. During the social novelty phase, young degus showed significantly higher sniffing-zone investigation frequency toward the novel intruder than toward the familiar cagemate (p = 0.0045; Figure 4D). In contrast, old degus did not show a significant difference in investigation frequency between the novel intruder and familiar cagemate zones (p = 0.1360; Figure 4D). Consistently, the social novelty discrimination index was significantly lower in old degus compared with young degus (p = 0.0065; Figure 4E). Using a social novelty discrimination index ≤ 0 to define reduced social novelty preference, 13 of 23 old degus (56.5%) showed reduced social novelty preference, compared with 1 of 11 young degus (9.1%; Fisher’s exact test, p = 0.011; Figure 4F). Together, these findings suggest that aging in laboratory-housed degus is associated with reduced novelty-directed social investigation, while baseline sniffing-zone preference and general sociability were not significantly different between age groups.

**Figure 4.**
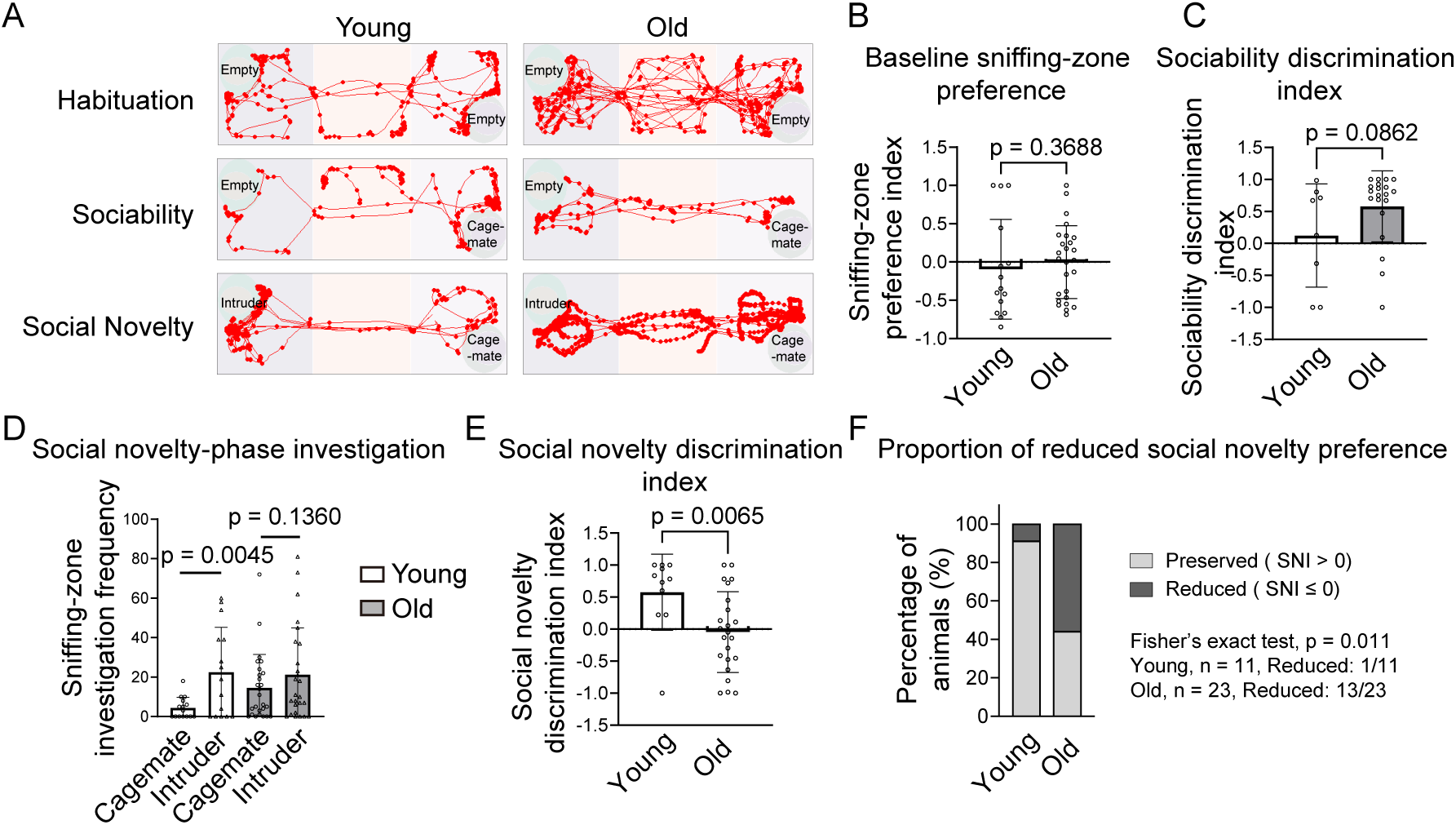
Aged degus show reduced social novelty preference. (A) Representative locomotor trajectories from young and old degus during the three-chamber social behavior assay, including habituation, sociability, and social novelty phases. During the sociability phase, a familiar cagemate was placed in one wire enclosure and the opposite enclosure remained empty. During the social novelty phase, a novel intruder was introduced into the previously empty enclosure, while the familiar cagemate remained in the opposite enclosure. (B) Quantification of baseline sniffing-zone preference during the habituation phase. The sniffing-zone preference index was calculated using the two enclosure-associated sniffing zones before introduction of social stimuli. (C) Quantification of sociability during the sociability phase. The sociability discrimination index was calculated as (cagemate − empty enclosure) / (cagemate + empty enclosure) using sniffing-zone investigation time. (D) Quantification of sniffing-zone investigation frequency toward the familiar cagemate and novel intruder during the social novelty phase. Young degus showed significantly higher investigation frequency toward the novel intruder than toward the familiar cagemate, whereas old degus did not show a significant difference between the two social stimuli. (E) Quantification of the social novelty discrimination index calculated as (novel intruder − familiar cagemate) / (novel intruder + familiar cagemate) using sniffing-zone investigation time. (F) Proportion of animals with reduced social novelty preference, defined as a social novelty discrimination index ≤0. Data are presented as mean ± SD with individual animals overlaid where applicable. Paired comparisons between familiar cagemate and novel intruder investigation frequency were performed using paired two-tailed t tests. Young and old index values were compared using two-tailed Mann–Whitney tests. Proportions were compared using Fisher’s exact test.

### 3.5. Sex-stratified analysis of open-field and social novelty behavioral metrics

Because both male and female animals were included in the behavioral cohort, we next examined whether the major behavioral metrics differed by sex within each age group. In the open-field arena, young males and young females did not differ significantly in total distance moved, mean velocity, moving frequency, cumulative movement duration, center-zone cumulative duration, or area-corrected thigmotaxis score (Figure 5A – F). Similarly, old males and old females did not show significant differences in these open-field locomotor or spatial-use metrics. We also examined the social novelty discrimination index in males and females within each age group. Social novelty discrimination index did not differ significantly between males and females within either the young or old group (Figure 5G). These findings indicate that, within this cohort, the analyzed open-field and social novelty behavioral metrics were not strongly influenced by sex within age groups. Together with the age-group comparisons shown above, these analyses suggest that the age-associated behavioral differences observed in old degus were not driven by a single sex-specific subgroup.

**Figure 5.**
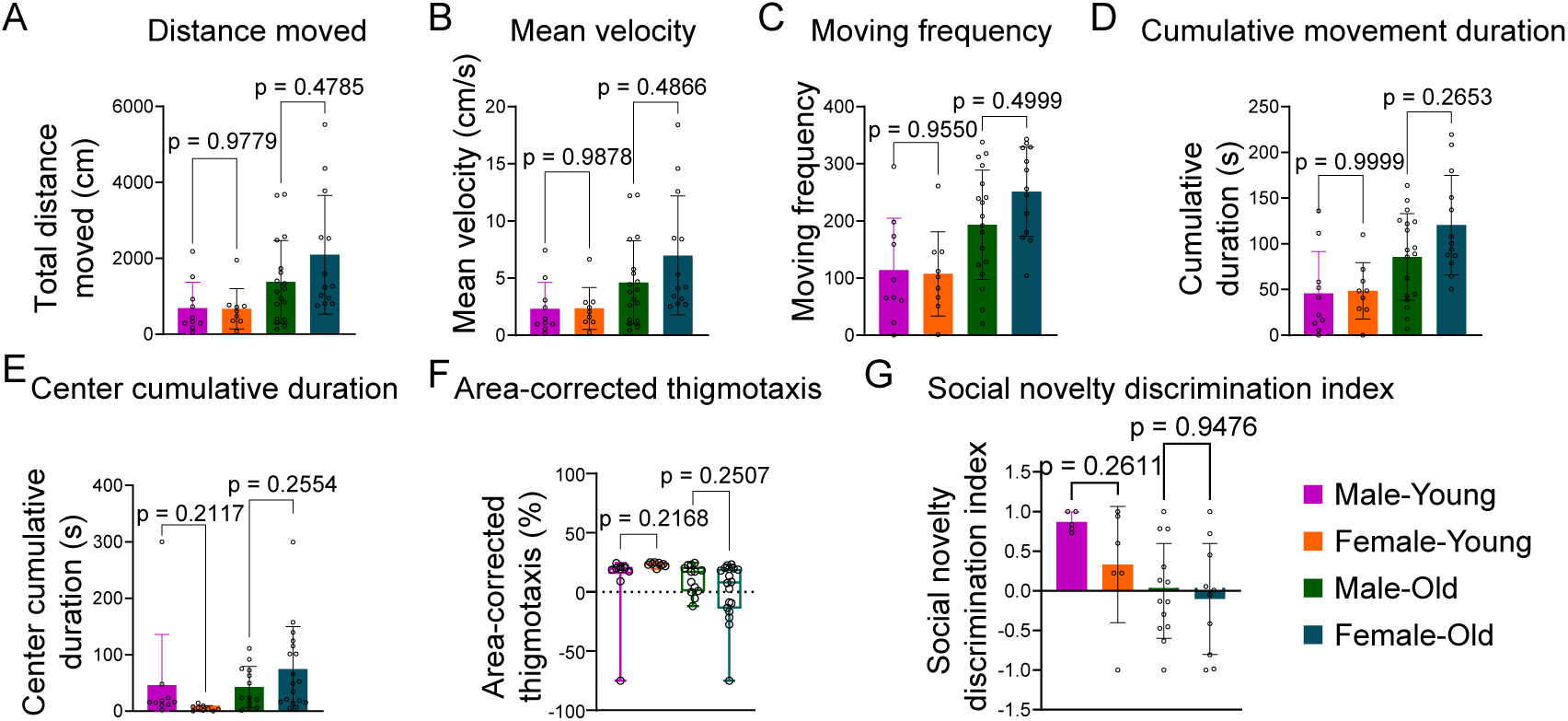
Sex-stratified analysis of open-field and social novelty behavioral metrics within young and old degus. (A – F) Sex-stratified quantification of open-field behavioral metrics, including total distance moved (A), mean velocity (B), moving frequency (C), cumulative movement duration (D), center-zone cumulative duration (E), and area-corrected thigmotaxis score (F). Open-field analysis included young males, n = 10; young females, n = 9; old males, n = 13; and old females, n = 18. (G) Sex-stratified quantification of the social novelty discrimination index. Social novelty analysis included young males, n = 7; young females, n = 8; old males, n = 10; and old females, n = 16. Within both young and old groups, males and females did not show significant differences in the analyzed open-field or social novelty metrics. Data are presented as mean ± SD with individual animals overlaid. Multiple comparisons were performed using Šídák’s multiple comparisons test.

### 3.6. Manual ethological scoring reveals increased rearing and fecal boli in aged degus

To complement automated behavioral tracking, we manually scored selected ethological behaviors during a 5-min open-field exposure. This analysis captured behavioral features that are not fully represented by distance- or velocity-based tracking metrics, including fecal boli, rearing, grooming, and jumping. Old degus produced significantly more fecal boli than young degus (p = 0.0020; Figure 6A) and showed a marked increase in rearing events (p < 0.0001; Figure 6B). Grooming events did not differ significantly between age groups (p = 0.1101; Figure 6C), whereas jumping events showed a non-significant trend toward increase in old animals (p = 0.0661; Figure 6D). Together, these manually scored behaviors indicate that aged degus exhibit increased fecal output and vertical exploration during open-field exposure, supporting an age-associated shift in stress-related and exploratory behavioral responses beyond automated locomotor measures.

**Figure 6.**
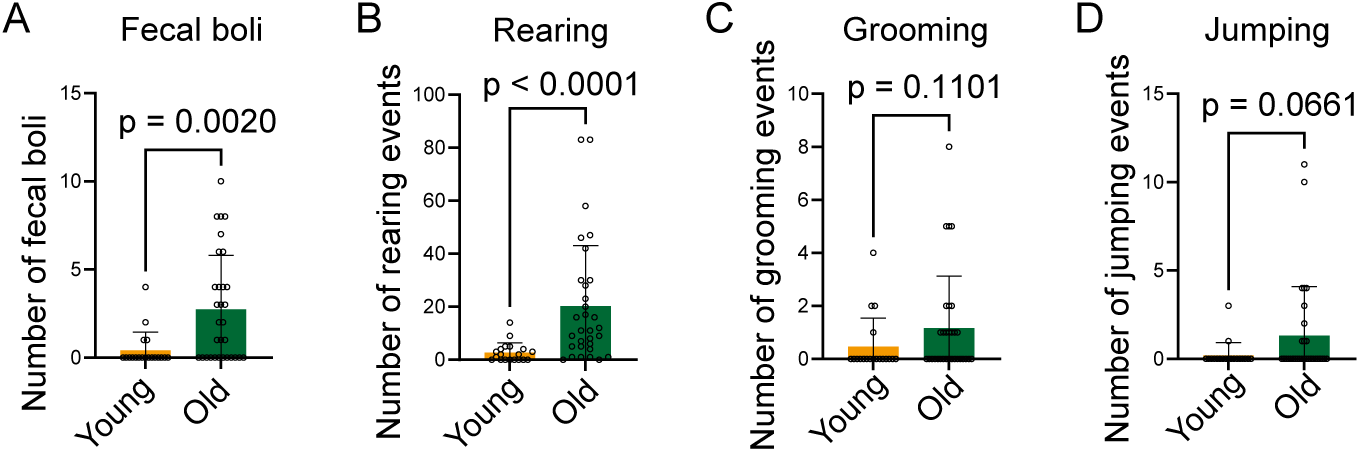
Manual ethological scoring of open-field behaviors in young and old degus. (A – D) Quantification of manually scored open-field behaviors during the 5-min open-field exposure, including fecal boli (A), rearing events (B), grooming events (C), and jumping events (D). Old degus showed significantly increased fecal boli and rearing events compared with young degus. Grooming did not differ significantly between age groups, whereas jumping showed a non-significant trend toward increase in old animals. Data are presented as mean ± SD with individual animals overlaid; young, n = 19 (10 males and 9 females); old, n = 31 (13 males and 18 females). Statistical significance was determined using a two-tailed Mann–Whitney test.

## 4. Discussion

This study reveals age-associated behavioral changes across locomotor, exploratory, stress-related, and social novelty domains in laboratory-housed Octodon degus. Rather than showing a generalized reduction in behavioral engagement, aged degus displayed a mixed phenotype characterized by increased locomotor activity during open-field and chamber-based testing, increased rearing and fecal output, reduced thigmotaxis, and reduced social novelty preference. Together, these findings indicate that aging in degus is associated with coordinated changes in exploratory activation, stress-related responses, center avoidance, and novelty-directed social investigation.

A prominent finding was that aged degus did not exhibit reduced locomotor engagement during behavioral testing. Old animals moved greater distances, showed higher velocity, had more movement bouts, and spent more time moving in both the open-field arena and chamber-based apparatus. This distinction is important because apparent deficits in social or cognitive assays can arise from reduced locomotor activity, limited task engagement, or insufficient exploration of the testing apparatus^18,32^. In the present study, reduced social novelty preference occurred despite increased locomotor activity, suggesting that this phenotype was not simply explained by reduced movement or general behavioral inactivity. However, because activity was measured during behavioral testing rather than in the home cage, these results should be interpreted as increased behavioral activation in novel testing environments, not as increased overall daily activity.

The open-field phenotype observed in aged degus was not consistent with a simple increase or decrease in anxiety-like behavior. Old animals spent more time in the center zone and showed reduced thigmotaxis, a pattern that may be consistent with reduced center avoidance. However, center-zone exploration is not a pure measure of anxiety-like behavior and can also be influenced by locomotor activity, exploratory drive, novelty response, and risk assessment^33–36^. In this study, reduced thigmotaxis occurred together with increased locomotor activity and rearing, suggesting that increased center-zone duration may partly reflect enhanced novelty-driven exploration rather than simply reduced anxiety-like behavior. At the same time, aged degus produced more fecal boli, a commonly used measure of emotionality- or stress-related autonomic output in open-field testing. Thus, the combination of increased exploratory activity, increased rearing, increased fecal output, and reduced thigmotaxis suggests altered arousal and stress responsiveness during exposure to a novel environment.

The reduction in social novelty preference provides a more direct link to cognitive and social-recognition-related behavioral domains. Degus are highly social animals, and successful performance in the three-chamber social novelty test requires animals to distinguish a novel conspecific from a familiar one and to direct investigation toward the novel social stimulus. Young degus showed greater investigation of the novel intruder than the familiar cagemate, whereas old degus did not show a significant novelty-directed investigation pattern. Consistently, aged degus had a lower social novelty discrimination index and a larger proportion of animals with reduced social novelty discrimination index. Because baseline sniffing-zone preference and sociability index were not significantly different between age groups, this phenotype is less likely to reflect pre-existing zone bias or a general loss of social approach. Instead, it may reflect reduced social novelty discrimination, altered social recognition memory, reduced attention to social cues, or changes in novelty-directed social motivation.

These behavioral changes may be relevant to neurodegenerative aging because AD and related dementias are associated not only with memory impairment, but also with alterations in affective behavior, arousal regulation, stress responsiveness, and social function^16,37–43^. In human AD, neuropsychiatric symptoms such as anxiety, agitation, altered stress responsiveness, and behavioral disinhibition are common and have been linked to disease progression and dysfunction of hippocampal, cortical, amygdalar, and limbic circuits^44–48^. Consistent with this broader clinical spectrum, behavioral phenotypes in AD mouse models are also heterogeneous and can include altered locomotor activity, anxiety-like behavior, exploration, stress responsiveness, social interaction, and novelty recognition. Importantly, the direction of these phenotypes can vary depending on genotype, age, sex, genetic background, disease stage, and testing paradigm^17,18,49–51^. Thus, open-field and social behavioral changes in aged animals should not be interpreted only as changes in motor output or memory, but as part of broader alterations in arousal, affective, exploratory, and social behavioral domains.

Previous degu studies have reported age-associated impairments in burrowing, recognition memory, social memory, and maze-based cognitive performance, in some cases together with AD-like neuropathological features^52–57^. Our findings extend this work by identifying coordinated changes in exploratory activation, stress-related open-field responses, and social novelty preference within the same laboratory-housed aging cohort. These findings should also be interpreted in light of the broader AD mouse-model literature, where open-field and anxiety-related phenotypes are highly model- and context-dependent. Across AD mouse models, locomotor and exploratory phenotypes can range from hypoactivity to hyperactivity, and anxiety-like measures can vary in direction depending on genotype, age, sex, genetic background, disease stage, and testing paradigm^58–66^. Thus, the increased locomotor activity and reduced thigmotaxis observed in aged degus should not be interpreted as preserved behavioral function or simply reduced anxiety. Instead, they support the interpretation that aging in degus is associated with altered behavioral activation and anxiety-related exploration, together with reduced novelty-directed social investigation. Rather than suggesting that aged degus phenocopy a specific AD mouse model, these results define a domain-specific behavioral profile that can be used in future studies to link natural degu aging with molecular, endocrine, and neuropathological markers.

Several limitations should be considered. This study was designed to define age-associated behavioral phenotypes and generate hypotheses about their neurobiological significance, rather than to identify causal mechanisms. Locomotor activity was not measured in the home cage, and open-field center-zone exploration and thigmotaxis do not directly define anxiety state. Social novelty behavior may also be influenced by multiple processes, including social recognition, novelty preference, olfaction, attention, motivation, and exploratory drive. In addition, animals with zero cumulative duration in both social sniffing zones were excluded from index-based analyses because a discrimination index could not be calculated. Finally, this study did not include neuropathological, endocrine, or molecular measurements in the same animals. Direct correlations with amyloid, tau, neuroinflammation, corticosterone, and other aging- or AD-related markers will be needed to determine whether these behavioral changes track neurodegenerative pathology or stress-system alterations in aged degus.

The coordinated alterations observed across exploratory, stress-related, and social novelty domains suggest that aged degus may exhibit broader changes in behavioral regulation that are relevant to brain aging and neurodegenerative vulnerability. Future studies linking these behavioral measures with amyloid pathology, tau phosphorylation, neuroinflammation, corticosterone signaling, and age-associated transcriptional or metabolic alterations will help determine whether they reflect neurodegenerative vulnerability, broader changes in arousal and stress responsiveness, or both. Such analyses will be important for establishing how behavioral heterogeneity in naturally aging degus relates to biological mechanisms of brain aging and to phenotypes relevant to age-related neurodegenerative disease.

## 5. Conclusions

In conclusion, this study defines an age-associated behavioral profile in laboratory-housed Octodon degus across locomotor, exploratory, stress-related, and social novelty domains. Aged degus did not show a generalized reduction in behavioral engagement; instead, they displayed increased locomotor activity during open-field and chamber-based testing, increased rearing and fecal output, reduced thigmotaxis, and reduced lower social novelty discrimination index. This pattern suggests that aging in degus is associated with enhanced exploratory activation, altered stress-related autonomic responses, reduced center avoidance, and reduced novelty-directed social investigation. These behavioral differences were not explained by significant male–female differences within age groups for the major open-field and social novelty metrics examined. Together, these findings provide a functional behavioral framework for future studies linking degu aging phenotypes to neurodegenerative vulnerability, behavioral heterogeneity, and molecular or neuropathological features of brain aging.

## Author Contributions

Conceptualization, H.B., A.S. and V.G.; methodology, H.B. and Y.L.; investigation, H.B. and Y.L.; formal analysis, H.B.; data curation, H.B.; visualization, H.B.; writing—original draft preparation, H.B.; writing—review and editing, H.B., Y.L., A.S. and V.G.; supervision, A.S. and V.G.; project administration, A.S. and V.G.; funding acquisition, A.S. and V.G. All authors have read and agreed to the published version of the manuscript.

## Funding

This research was funded by US National Institutes of Health AG047200 grant to VG and AS.

## Informed Consent Statement

Not applicable.

## Data Availability Statement

The data presented in this study are available within the article. Additional data supporting the findings of this study are available from the corresponding author upon reasonable request.

## Acknowledgments

The authors thank members of the Gorbunova and Seluanov laboratories for helpful discussions and support with degu colony maintenance and behavioral studies. The authors also thank the animal care staff at the University of Rochester for assistance with animal husbandry.

## Conflicts of Interest

The authors declare no conflicts of interest.

## Notes

### Competing Interest Statement

The authors have declared no competing interest.

